# Spatial organization of the transcriptional regulatory network of *Saccharomyces cerevisiae*

**DOI:** 10.1101/509224

**Authors:** Dong-Qing Sun, Liu Tian, Bin-Guang Ma

## Abstract

Transcriptional regulatory network (TRN) is a directed complex network composed of all regulatory interactions between transcription factors and corresponding target genes. Recently, the three-dimensional (3D) genomics studies have shown that the 3D structure of the genome makes a difference to the regulation of gene transcription, which provides us with a novel perspective. In this study, we constructed the TRN of the budding yeast *Saccharomyces cerevisiae* and placed it in the context of 3D genome model. We analyzed the spatial organization of the yeast TRN on four levels: global feature, central nodes, hierarchical structure and network motifs. Our results suggested that the TRN of *S. cerevisiae* presents an optimized structure in space to adapt to functional requirement.

## Introduction

A transcription factor (TF) is a protein that controls the rate of the production of messenger RNA by binding to transcription factor binding site of a DNA template, which can regulate the expression of genes in response to signals inside or outside the cell. So far, nearly 300 TFs have been found in the model eukaryote *Saccharomyces cerevisiae*, which function as activators or repressors to regulate the expression of more than 5000 genes in the genome. Genes regulated by transcription factors are referred to as target genes (TGs). A transcription factor can regulate one or more target genes, and a gene can be regulated by multiple transcription factors. Transcription factors themselves are proteins encoded by genes, and these coding genes are also regulated by other TFs. In this way, all the regulatory interactions linking transcription factors to their corresponding target genes are built into the transcriptional regulatory network (TRN). The existence of TRN helps to coordinate the gene expression on a genomic scale. In TRN, a node represents a gene and an edge manifests the relationship in which a gene (TG) is regulated by the protein products of another gene (TF).

TRN, modeled as a directed graph, has its characteristic topology, the global and local properties of which have been widely studied [1-4]. At the global level, TRN displays a scale-free feature with a power-law degree distribution, which means some genes are architecturally significant and thus functionally important. Besides, Yu *et al.* identified pyramid-shaped hierarchical structures in the TRNs of *Escherichia coli* and *Saccharomyces cerevisiae* and found that genes in different levels of hierarchy take on different responsibilities [2]. With respect to the TRN of *S. cerevisiae*, dynamic and evolutionary properties of hierarchies have also been investigated [5,6]. At the local level, network motifs in TRN are considered significant. Network motifs are connected patterns that occur more frequently in the real network than expected in random networks. Motifs can be found in many complex networks, such as social networks, protein-protein interaction networks and regulatory networks. Motifs, as the building blocks of TRN, have important dynamic functions [7]. For example, feed-forward loop (FFL) plays a role in filtering out spurious pulses of signal and accelerating response, while single input module (SIM) can regulate the expression of a group of genes with shared function in a temporal order [8].

Although genome annotations of species become increasingly perfect, how the discrete regulatory elements interact in space to precisely control the expression of genes is still unclear. It has started to explore the spatial organization of genes and their regulatory elements. In 2002, Dekker and co-workers first developed the chromosome conformation capture (3C) technique to infer the physical proximity of a single pair of genomic loci, which is based on the premise that the physical proximity of chromatin is associated with the contact frequency between DNA segments in cell populations *in vivo* [9]. Since then, 3C-derived methods emerged successively, including 4C, [10], 5C [11], Hi-C [12] and ChIA-PET [13]. With these techniques, 3D chromosome models of many eukaryotic organisms [14-16] have been obtained and the yeast is one of them [14].

The chromatin contact data generated by 3C-based methods has allowed further understanding of the relationship between structure and function, including the idea that long-range associations are a common control mechanism of gene expression, which play an important role in determining the nuclear organization [17]. In addition, sub-nuclear position preference of chromosomes in the human genome has been uncovered by Hi-C [12]. Recently, Tokuda and Sasai analyzed the difference of genome structures between the mutants and the wide-type of yeast, and found that the transcriptional activity of genes is related to their spatial distribution inside the nucleus [18]. These findings imply that genes regulated by their TFs are not randomly distributed in 3D space and that for TRN there must be certain spatial organization to maintain the high efficiency of transcriptional regulation. In order to achieve a comprehensive understanding of the spatial organization of the yeast TRN, we mapped the constructed TRN into the 3D genome model of *S. cerevisiae* and then analyzed it on both global and local levels, including regulation directions, central nodes, hierarchical structure and network motifs.

## Materials and Methods

### Constructing the yeast TRN

The transcriptional regulatory network of *S. cerevisiae* was derived from SGD database [19]. The network consists of 186 transcription factors and 5727 target genes, which involve 28260 regulatory interactions.

### Determining central nodes

The topology of the yeast TRN exhibits scare-free property, which indicates a fraction of nodes have high degrees, while most nodes only participate in few connections. We calculated the in-degree centrality, out-degree centrality, betweenness centrality and closeness centrality for all the genes in the yeast TRN, and accordingly we defined the genes with top 5% highest values of in-degree centrality as in-hubs and the genes with top 15% highest values of out-degree centrality, betweenness centrality and closeness centrality as out-hubs, bottlenecks, centers, respectively. All the rest genes were defined as others.

### Determining the spatial distance of regulatory interaction

The 3D model of the yeast genome was derived from the study of Duan *et al.* [14], which provides the 3D coordinates of genomic loci. We mapped all the genes that participate in transcriptional regulation to the 3D genomic model and calculated the spatial distance (Euclidean distance in nm) between the transcription factor genes and corresponding target genes.

### Determining the hierarchical levels of the yeast TRN

We divided the yeast TRN into four hierarchical levels using the method adopted by Kumar *et al.* [20]. Firstly, genes without outgoing edges in the directed graph were classified as target genes (TGs) which comprise the lowest level in the hierarchy: Target level. These target genes with their edges were then excluded from the TRN so that we obtained a subgraph of the full TRN, which comprises only transcription factors and their regulatory interactions. Next, we identified all the strongly connected components (SCCs) in the subgraph and collapsed every SCC into a super node. SCC is a strongly connected subgraph in which there is a path in each direction between each pair of nodes. Actually, self-loops belong to SCC. And then we substituted the edges associated with (to or from) the genes in each SCC for the edges to (or from) the corresponding super node, whence the original subgraph was turned into a directed acyclic graph (DAG). The transcription factors in the DAG were categorized into three levels: nodes with only outgoing edges were classified into the Top level, nodes with both outgoing and incoming edges were classified into the Middle level, and the rest nodes were classified into the Bottom level. Finally, the whole TRN was decomposed into four hierarchical levels (Top, Middle, Bottom and Target).

### Finding network motifs

Mfinder is one of the tools for network motif detection [21]. We identified 3-nodes motifs with Mfinder and found 15801 feed forward loops (FFLs); 56 single input modules (SIMs, in which the only one TF regulates at least two TGs) in the full yeast TRN were identified with an in-house Python program according the definition from a previous paper [22].

## Results

We built the transcriptional regulatory network of budding yeast with the regulation data from SGD database [19], which includes 5913 nodes (genes) and 28260 edges (regulatory interactions) (Supplementary Data). Based on the 3D model of the yeast genome [14], we analyzed the spatial structure of the yeast TRN from four aspects including global features, central nodes, hierarchical structure and network motifs. With the influence of the linear proximity in the genome taken into consideration, we repeated the above analysis considering only regulations of which TF genes and target genes are separated by at least 50 kb in genome, and we found that whether close-range regulatory interactions were excluded or not did not change the results. The results presented in the article are from the analysis of the whole TRN.

### Global features

We defined the spatial distance of a regulatory interaction as the Euclidean distance between the TF gene and target gene in the 3D genome model. We calculated the distance of each regulatory interaction in the whole TRN. Likewise, we calculated the distance between any two genes in the entire genome. According to the result of statistical test, we found that the spatial distances between gene pairs with regulatory interaction (named as TRN, Table S1, average: 82.86nm) are significantly (Mann-Whitney-Wilcoxon test, *P* < 2.2e-16) longer than those among all genes in the genome (average: 80.52nm). According to the direction of regulation, regulatory interactions can be divided into four types: positive (+, activation), negative (-, repression), bidirectional (+/-) and unknown (?/binding enriched) (Supplementary Data). We compared the spatial distances among regulatory interactions with different directions. We found that the distances of positive regulations are comparable with those of the whole TRN, while the distances of negative regulations are significantly longer (Figure 1B and Table S1, S2).

**Figure 1.**
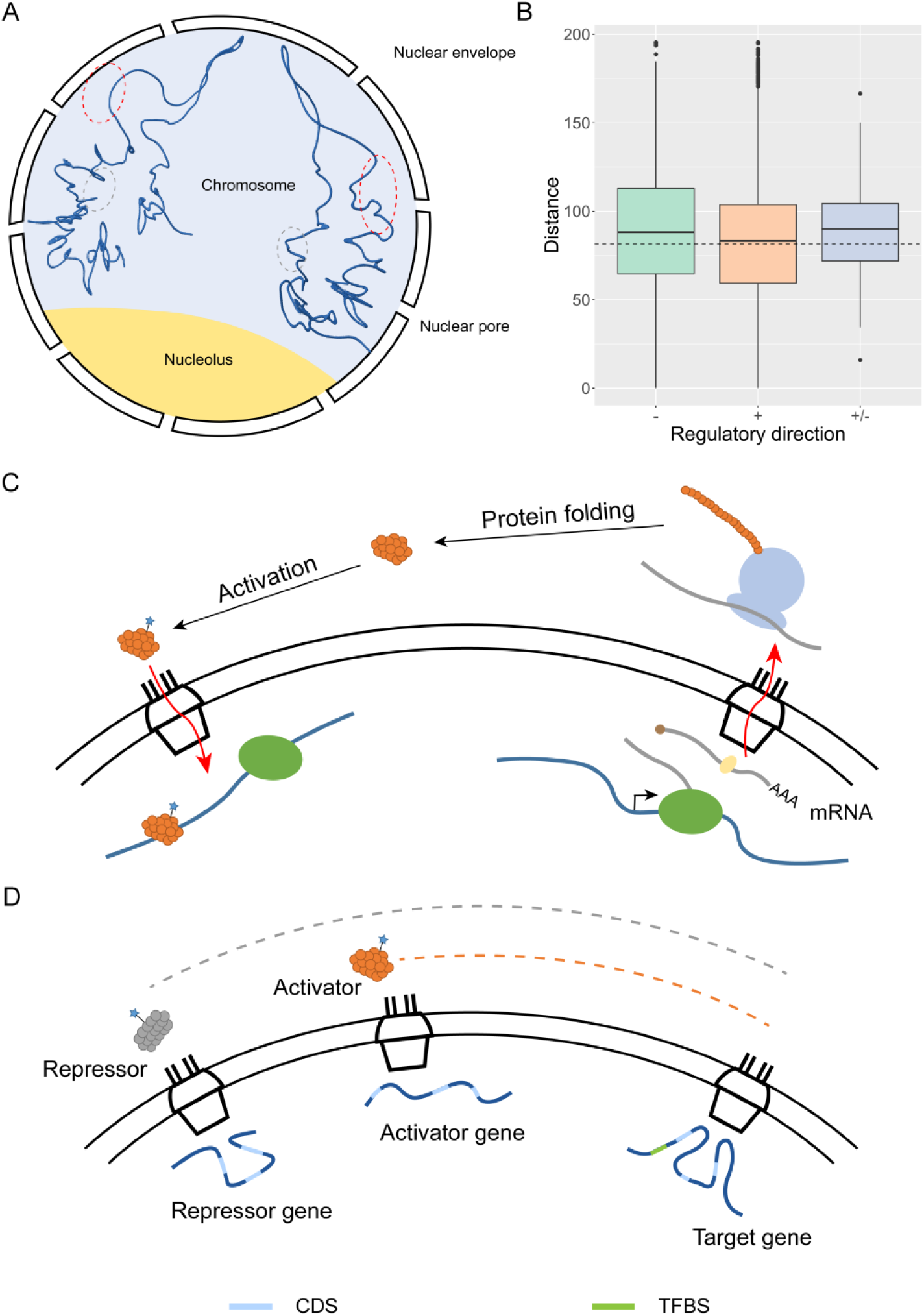
The global spatial organization of yeast TRN. (A) Genes involving transcriptional regulation tend to be located near the nuclear envelope (the area within the red dashed oval), while genes located near the center of the nucleus (the area within the grey dashed oval) are unlikely to participate in transcription. (B) The distribution of spatial distances of different regulatory directions is shown in box plot. The black dashed line indicates the median distance of all regulations in the TRN. (C) The schematic diagram illustrates the process of TF generation and the following transcriptional regulation it involves in physical space. (D) As shown, a shorter distance between the regulator-coding gene and its target gene might allow early access to the target gene. If the activator-coding gene is closer to its target gene, it will be beneficial to the efficiency of regulation.

The results indicate the spatial distribution of genes in the cell nucleus that genes involving transcriptional regulation are located near the nuclear envelope (Figure 1A). As eukaryote, *S. cerevisiae* performs its transcription and translation in the nucleus and cytoplasm, respectively. After a TF is generated, it needs to enter the nucleus to regulate its target genes (Figure 1C). The locations of the genes at the nuclear periphery may benefit them to be easily regulated. As for the difference between positive and negative regulations, it seems that the TFs involving negative regulations have to travel a longer distance to repress their target genes, whereas the TFs involving positive regulations can reach their target genes through a shorter distance (Figure 1D). Such a distribution matches their respective features that negative regulations are supposed to be secure and positive regulations are supposed to be efficient.

### Central nodes

We sought to explore the spatial distribution of four categories of central nodes: in-hubs, out-hubs, bottlenecks and centers. The in-hubs are those highly regulated genes which may encode important proteins. The statistical test results show that the distances of the regulatory interactions that in-hubs participate in (both in-hub to in-hub and in-hub to others) are significantly shorter than those of the whole TRN (Table S3), indicating that the in-hubs are co-localized with their partners in space. The distances of regulatory interactions between out-hubs and others are significantly longer than those of the TRN (Table S3), indicating that out-hubs who have more target genes disperse in space so that they could regulate their widespread target genes.

Bottlenecks are defined according to betweenness centrality and denote the nodes controlling information flow in the network, deletion of which may lead to cell death [3]. In TRN, bottlenecks are the TFs that many other TFs must regulate in order to regulate their real target genes. The distances of regulatory interactions between bottlenecks and others are significantly longer than those of the TRN (Table S3), indicating that bottlenecks are distributed separately, which will be beneficial to massive regulation cascades that pass through bottlenecks.

Centers are defined according to closeness centrality and denote the nodes with shortest paths to other reachable nodes. The distances of regulatory interactions that only center TFs participate in are similar to those of the TRN, but the distances between centers and other genes are significantly longer than those of the TRN (Table S3), indicating that center TFs are distributed evenly in the 3D space, each of which plays an import role within a local region and once beyond the range that a local center TF can work, the distances between many other genes and this TF get longer.

### Hierarchical structure

All the genes were classified into four levels: Top, Middle, Bottom and Target. The genes in the Top level are transcription factors which only regulate genes but are not regulated by other TFs. The Middle regulators regulate genes and are also regulated by other regulators. The Bottom regulators do not regulate other TF genes but they regulate other non-TF target genes. The Bottom genes many also be regulated by the Top level TFs. What belong to the Target level are those non-TF genes mainly encoding enzymes related to metabolism, and all the TFs in the Top, Middle and Bottom levels can regulate the Target level genes. Indeed, the yeast TRN presents a pyramid-shaped hierarchical structure [2] (Figure S1 and Table S4). We counted the regulatory interactions of and between different levels (Table S5). As shown (Table S4 and Table S5), we there are 81 genes in the Middle level, which are involved in 24608 interactions with the Target level genes. Besides, the Middle level genes connect genes in other levels. The high connectivity of the Middle level TFs may indicate they play an import role in passing information and regulating expression of genes in response to stimuli.

According to the hierarchy of TRN, all regulations were divided into 8 types: Top-Middle, Top-Bottom, Top-Target, Middle-Middle, Middle-Bottom, Middle-Target, Bottom-Bottom and Bottom-Target. We compared the average spatial distances of each regulation type with that of the whole TRN (Figure 2A and Table S6). The results show that the spatial distances of the three regulation types that the Top level involves are extremely significantly shorter than the average distance of the whole TRN, while the distances of Middle-Target type are the opposite. The spatial distances of Middle-Middle, Middle-Bottom and Bottom-Target types show no significant difference with those of the whole TRN (Table S6).

**Figure 2.**
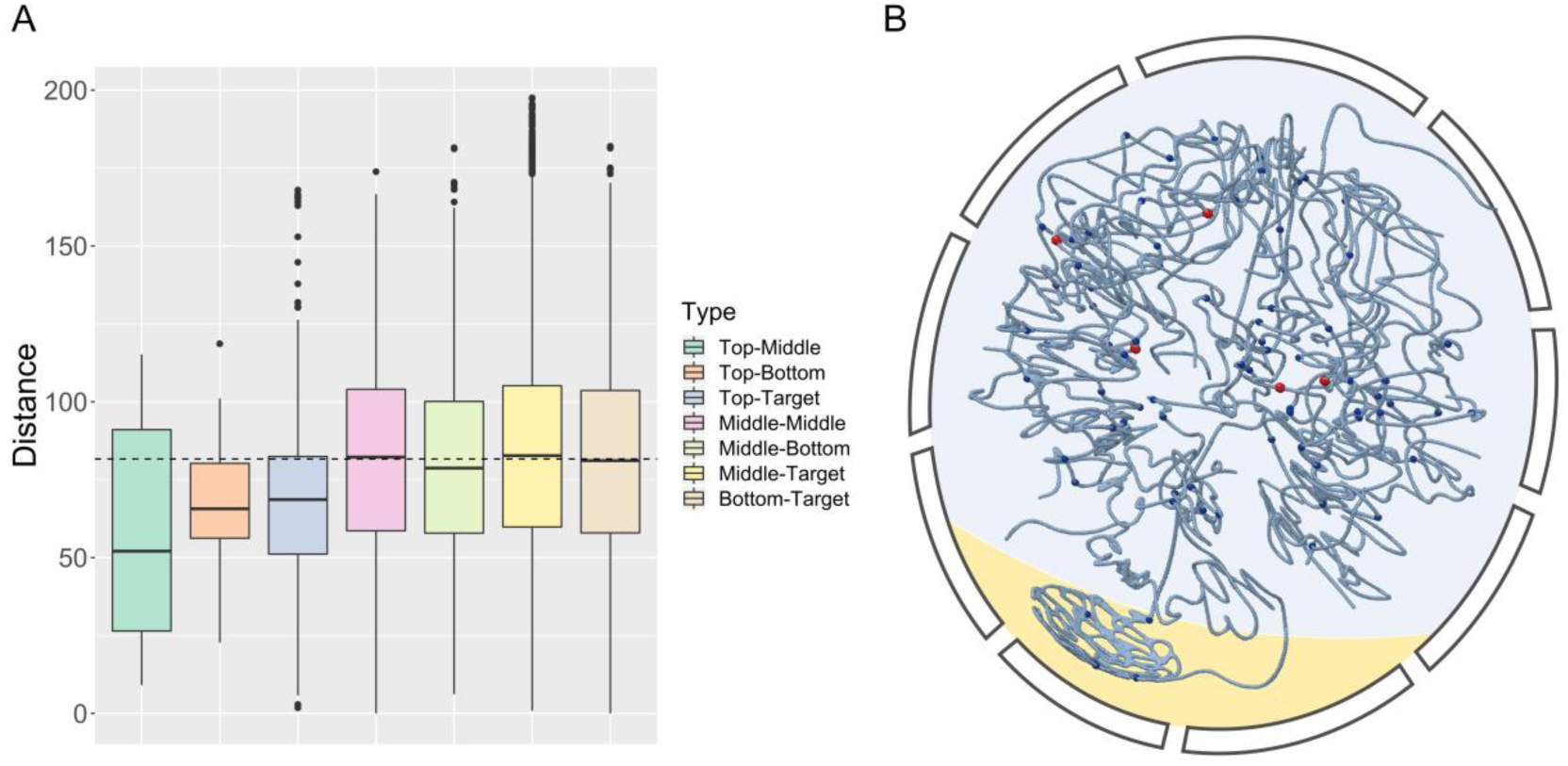
The spatial organization of the network hierarchy. (A) The box plot depicts the distribution of the spatial distances of regulations between different levels of hierarchy in the yeast TRN. The black dashed line indicates the median distance of all the regulations in the whole TRN. (B) Top-level and middle-level genes are shown as spheres in the 3D model of the yeast genome. Top-level genes (red) are mainly located near the middle of the chromosome arms, while middle-level genes (blue) are distributed evenly in the 3D genome.

Genes of the four levels were visualized in the 3D genome model (Figure 2B) by using RasMol [23]. We discovered that most of the Top level genes are located in the center of the nucleus. If the 3D genome is regarded as a sphere, the Top level genes roughly lie in the plane that passes through the center point of the sphere. In this way, it is not difficult to explain why the spatial distances of the regulation types that the Top level genes involve are smaller compared with the median distance of the entire network. It was reported that the Top level TFs usually participate in the initiation steps of biological processes [2]. Such a distribution of the Top level genes in 3D space will make initiation processes more efficiently. By comparison, it is also reasonable that the Middle level genes, which function as the main executors of transcriptional regulation, are evenly distributed throughout the 3D genome.

### Network motifs

Network motifs are defined as overrepresented connection patterns or sub-graphs in the natural networks compared with randomized networks. In TRNs, feed-forward loop (FFL), in which a master TF regulates a secondary TF and they both regulate a target gene, has been widely studied due to its functional importance and three-node simplicity. According to the direction of the three regulations, FFLs can be classified into eight types [24]. The coherent type-1 FFL (C1-FFL) occurs most frequently, followed by the incoherent type-1 FFL (I1-FFL), and the other six types of FFL occur much less frequently [25]. C1-FFL (Figure 3A), as a sign-sensitive delay element, can filter out the brief fluctuations in input stimuli, which helps to protect the target genes from noise interference [24]. The incoherent type-1 FFL (I1-FFL) (Figure 3A) can serve as a pulse generator and a response accelerator [26]. Single-input module (SIM) is another important motif in the TRN. In a SIM (Figure 3A), the sole regulator regulates a group of target genes and these target genes are not regulated by other regulators. Such a pattern helps SIM to coordinate the expression of a group of functionally related genes in a temporal order [8]. In this article, we selected FFL and SIM to study the spatial distribution of network motifs.

**Figure 3.**
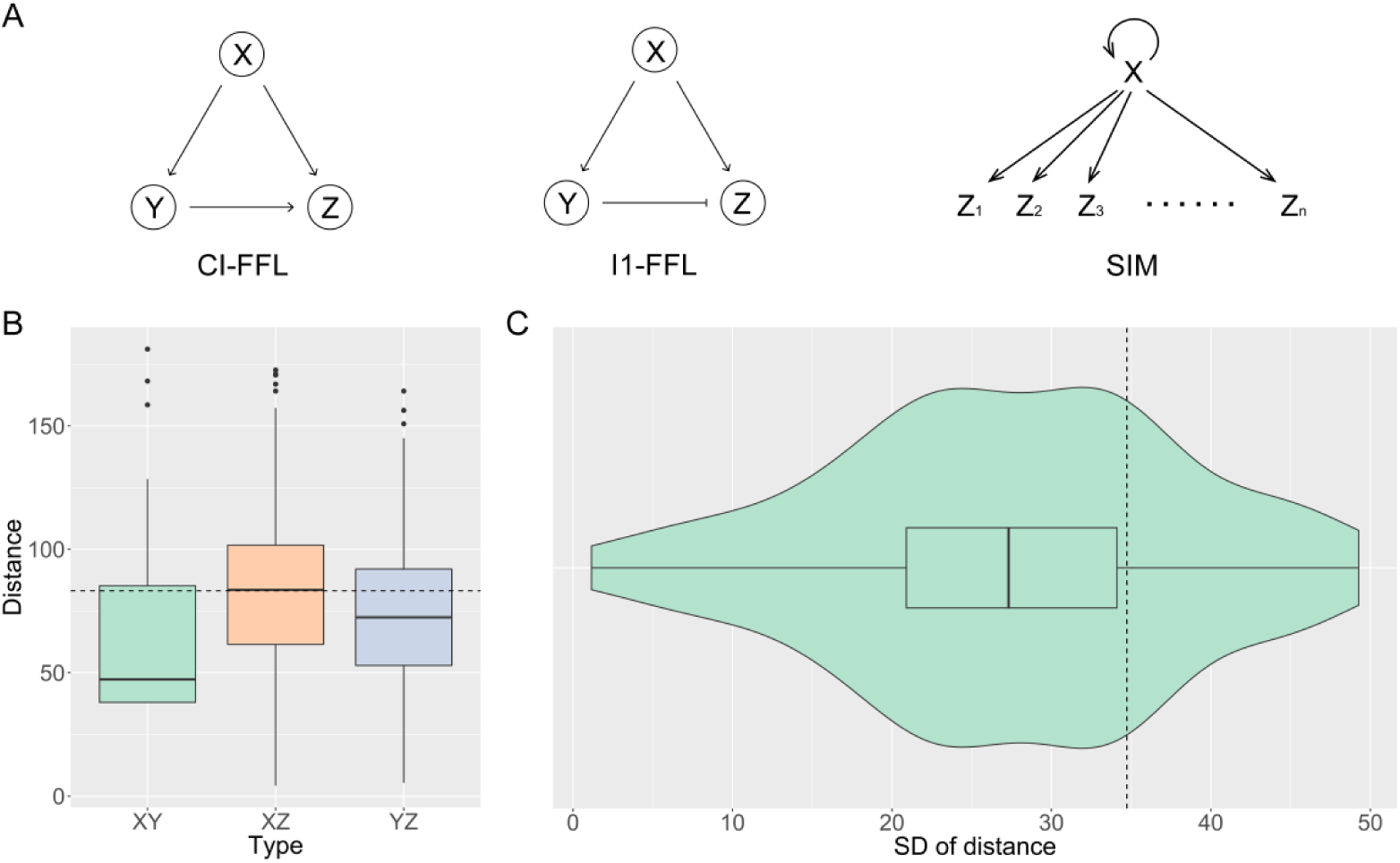
The spatial organization of network motifs. (A) The left and the middle show the regulation mode of CI-FFL and I1-FFL, respectively. The right shows the regulation mode of SIM, in which transcription factor X regulates a group of genes Z_1_, Z_2_, Z_3_, …, Z_n_, and X tends to regulates itself. (B) The box plot depicts the distribution of the spatial distances of regulation edges within C1-FFLs. The black dashed line indicates the median distance of all the positive regulations in the TRN. (C) The distribution of the standard deviation (SD) of distances within SIMs is shown with the violin plot and box plot. The black dashed line indicates the standard deviation of distances of all the regulations in the whole yeast TRN.

We analyzed the spatial distances of all the FFLs and SIMs. The distance of a FFL (SIM) is determined as the average spatial distance of all regulatory interactions within a FFL (SIM). The spatial distances within FFLs are statistically compared with those of the whole TRN by Mann-Whitney-Wilcoxon test. We found that the spatial distances within FFLs are significantly longer than the TRN and that the distances within C1-FFLs are significantly shorter than the TRN, while the distances within I1-FFLs is significantly longer than the TRN (Table S7). We analyzed the distances of three regulation edges within C1-FFLs in detail. In a C1-FFL (Figure 3B and Table S8), the master TF (X) activates the secondary TF (Y) and both of them activate the target gene (Z). We found that the spatial distance of XZ is longer than those of XY and YZ to a very significant level and are comparable with those of all the positive regulations in the TRN. Moreover, the distance of XY is significantly shorter than that of YZ and they are both significantly shorter than those of all the positive regulations in the TRN.

Surprisingly, the spatial distances within SIMs are statistically equivalent to those of the whole TRN. But after calculating the standard deviation (SD) of distances in each SIM and that of the whole TRN, we found that the SDs of most SIMs (42 of 56) are smaller than that of the whole TRN. The result reflects the lower dispersion of distances within SIMs (Figure 3C). In a SIM, the uniformity of distances from the main regulator to its target genes can reduce the impact of distance difference between TF and TGs on the required temporal order of gene activation, thereby ensuring the coordinated expression of genes.

## Discussion

The transcriptional regulatory network provides a framework that helps to elucidate the organizational principles of transcriptional regulation. The development of the chromosome conformation capture technologies enables us to recognize TRN from the spatial angle. Placing TRN in the context of 3D genome, we calculated the 3D spatial distances of all regulations in the TRN and analyzed the distances from both global and local viewpoints, including the whole TRN, positive and negative regulations, central nodes, hierarchical structure and network motifs.

Taken as a whole, the genes participating in transcriptional regulation are likely located near the nuclear envelope (Figure 1A). Intuitively, the location near the nuclear periphery will facilitate the export of mature mRNA to the cytoplasm and the access of transcription factors to the genome. The gene-gating hypothesis, that highly expressed genes are associated with the nuclear pore complexes (NPCs) by transient interactions between transcribed locus and the NPCs, can be used to explain the rationality of such a 3D gene distribution [27]. But it is hard to interpret our results from the perspective of chromosome territories (CTs), which stated that large and gene-poor chromosomes are enriched near the nuclear periphery, while small and gene-rich chromosomes tend to be located near the center of the nucleus [12]. Given that the existence of CTs in yeast is still controversial [28], further analysis should not be done for now. The distances of positive regulations are shorter compared with negative regulations, probably indicating that positive regulations are more economical and efficient in terms of spatial distribution.

From the local perspective, the spatial organization of the yeast TRN has been optimized. First, for central nodes, the separate distribution of out-hubs, bottlenecks and centers of the TRN in 3D space allows them to be involved in regulation of massive genes. In the hierarchical structure, the Top level TFs are located in the middle of the chromosome arms, namely, near the center of the whole 3D genome, which makes it possible to start a biological process efficiently. In addition, the spatial distribution within network motifs also reflects the optimization of the TRN. The difference of three regulation edges in C1-FFLs is manifested in that the distances of XY and YZ are shorter than that of XZ and that of all the positive regulations in TRN. The shorter distance of XY will be convenient for activating the promoter of Y quickly to start Y accumulation for co-regulating the expression of Z by both X and Y. The spatial distances of regulations within a SIM tend to be similar to each other, which benefits the coordinated expression of genes with shared function in a temporal order by excluding the possible influences of spatial distances.

## Conclusion

In this study, we combined the transcriptional regulatory network and the 3D genome model of *S. cerevisiae* and analyzed the spatial organization of the yeast TRN from four aspects: global features, central nodes, hierarchical structure and network motifs. The results indicate that the yeast TRN adopts an optimized 3D structure to adapt to the functions at different levels, which provides new insights into our understanding on gene regulation mechanisms in real 3D space.

## Authors’ contributions

MBG conceived and supervised the study. MBG, SDQ, TL designed the study. SDQ performed the experiments. SDQ, TL and MBG analyzed the results. SDQ and MBG wrote the manuscript. TL revised the manuscript.

## Competing interests

The authors have declared no competing interests.

## Acknowledgements

This work was supported by the National Natural Science Foundation of China (Grant 31570844), Project 2662016PY094 supported by the Fundamental Research Funds for the Central Universities. The funders had no role in study design, data collection and interpretation, or the decision to submit the work for publication.

## Abbreviations

TRN: transcriptional regulatory network
3D: three-dimensional
TF: transcription factor
TG: target gene
FFL: feed-forward loop
SIM: single-input module

