## Supplementary Materials for "Spatial organization of the transcriptional regulatory network of *Saccharomyces cerevisiae*"

**Table S1** Statistics of regulations of different directions in the yeast TRN

| Type of regulations | Number of regulations | Average spatial distance |
|---|---|---|
| + | 6114 | 83.42 |
| - | 3211 | 89.79 |
| +/- | 18 | 90.89 |
| TRN | 28260 | 82.86 |

Note: '+' represents positive regulation (i.e., activation), '-' represents negative regulation (i.e., repression) and '+/-' represents bidirectional regulation.

**Table S2** Mann-Whitney-Wilcoxon tests for the comparisons of spatial distances of regulatory interactions with different directions in the yeast TRN (see also **Table S1**)

| Type | P-value | Significance |
|---|---|---|
| TRN_+ < TRN_- | 8.09e-15 | ** |
| TRN_+ != TRN | 0.1679 | |
| TRN_- > TRN | < 2.2e-16 | ** |

Note: 'TRN_+' represents positive regulation (i.e., activation), 'TRN_-' represents negative regulation (i.e., repression) and 'TRN' represents all regulation in the TRN. '<', '!=' and '>' represent being significantly shorter, of no significant difference and significantly longer, respectively. '**' indicats the significance level of 0.01.

**Table S3** Mann-Whitney-Wilcoxon tests for the comparisons of the distances of regulatory interactions that central nodes involve in the yeast TRN

| Type | | P-value | Significance |
|---|---|---|---|
| In-hub | in-hub < TRN | 0.0087 | ** |
| | inter < TRN | 5.44e-05 | ** |
| Out-hub | out-hub != TRN | 0.3544 | |
| | inter > TRN | 0.0358 | * |
| Bottleneck | bottleneck != TRN | 0.8859 | |
| | inter > TRN | 1.95e-08 | ** |
| Center | center != TRN | 0.8305 | |
| | inter > TRN | 0.0256 | * |

Note: '<', '!=' and '>' represent being significantly shorter, of no significant difference and significantly longer, respectively. '**' indicates the significance level of 0.01 and '*' indicates the significance level of 0.05.

**Table S4** Number of nodes in different levels of hierarchy in the yeast TRN

| Hierarchy | Top | Middle | Bottom | Target | Total |
|---|---|---|---|---|---|
| Node number | 5 | 81 | 100 | 5727 | 5913 |

**Table S5** Statistics of regulations between different levels of hierarchy in the yeast TRN

| Type of regulation | Number of regulation | Average spatial distance |
|---|---|---|
| Top-Middle | 17 | 59.07 |
| Top-Bottom | 22 | 66.67 |
| Top-Target | 1148 | 67.45 |
| Middle-Middle | 318 | 81.10 |
| Middle-Bottom | 396 | 80.51 |
| Middle-Target | 24608 | 83.77 |
| Bottom-Bottom | 2 | 0.00 |
| Bottom-Target | 1749 | 81.47 |
| TRN | 28260 | 82.86 |

**Table S6** Mann-Whitney-Wilcoxon test for the comparisons of spatial distances between different levels of hierarchy with the whole yeast TRN (see also **Table S5**)

| Type | *P*-value | Significance |
|---|---|---|
| Top-Middle < TRN | 0.0079 | ** |
| Top-Bottom < TRN | 0.0092 | ** |
| Top-Target < TRN | < 2.2e-16 | ** |
| Middle-Middle != TRN | 0.7805 | |
| Middle-Bottom != TRN | 0.1381 | |
| Middle-Target > TRN | 0.0012 | ** |
| Bottom-Target != TRN | 0.2625 | |

Note: '<', '!=' and '>' represent being significantly shorter, of no significant difference and significantly longer, respectively. '**' indicates the significance level of 0.01.

**Table S7** Mann-Whitney-Wilcoxon test for the comparisons of the spatial distances within FFLs and those of the whole yeast TRN

| Type | P-value | Significance |
|---|---|---|
| FFL > TRN | < 2.2e-16 | ** |
| C1-FFL < TRN | 4.131e-11 | ** |
| I1-FFL > TRN | 0.02989 | * |

Note: ‘**’ indicates the significance level of 0.01 and ‘*’ indicates the significance level of 0.05.

**Table S8** Mann-Whitney-Wilcoxon test for the comparisons of the spatial distances of different regulatory edges within C1-FFL with those of the whole TRN

| Type | | P-value | Significance |
|---|---|---|---|
| C1-FFL | XZ > XY | < 2.2e-16 | ** |
| | XZ > YZ | 6.83e-07 | ** |
| | XY < YZ | 1.35e-11 | ** |
| | XY < TRN_+ | < 2.2e-16 | ** |
| | XZ != TRN_+ | 0.8905 | |
| | YZ < TRN_+ | 7.87e-10 | ** |

Note: ‘TRN_+’ represents positive regulations in the TRN. ‘<’ and ‘>’ represent being significantly shorter and longer, respectively. ‘**’ indicates the significance level of 0.01.

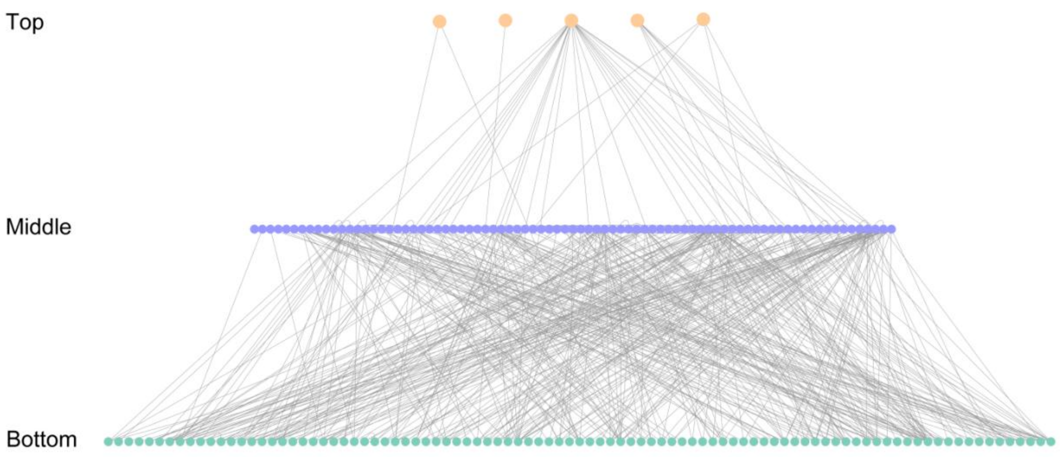


**Figure S1** The hierarchical structure of the yeast TRN. From the top down, Top (red), Middle (blue) and Bottom (green) TFs are organized in a hierarchical way. TGs are supposed to be below the Bottom TFs, but they are not displayed due to their large number.
